# Genomics of Egyptian Healthy Volunteers: The EHVol Study

**DOI:** 10.1101/680520

**Authors:** Yasmine Aguib, Mona Allouba, Alaa Afify, Sarah Halawa, Mohamed ElKhateb, Marina Sous, Aya Galal, Eslam Abdelrahman, Nairouz Shehata, Amr El Sawy, Mohamed Maghawry, Shehab Anwer, Omnia Kamel, Wessam El-Mozy, Hadir Khedr, Ahmed Essam, Nagwa Thabet, Pantazis Theotokis, Rachel Buchan, Risha Govind, Nicola Whiffin, Roddy Walsh, Heba Aguib, Ahmed Elguindy, Stuart A Cook, Paul J Barton, James Ware, Magdi Yacoub

**Author notes:** authors contributed equally (Yasmine Aguib, Mona Allouba).

## Abstract

Comprehensive genomic databases offer unprecedented opportunities towards effective tailored strategies for the prevention and treatment of disease. The integration of genomic and phenotypic data from diverse ethnic populations is also key to advancements in precision medicine and novel diagnostic technologies. Current reference genomic databases, however, are not representative of the global human population, making variant interpretation challenging and uncertain, especially in underrepresented populations such as the North African population. To address this, a study of 391 Egyptian healthy volunteers (EHVols) was initiated as a milestone towards establishing the 1000 Egyptian Genomes project.

## INTRODUCTION

Cardiovascular disease is a major cause of death and disability worldwide (Murray & Lopez, 2017; WHO 2017) and its prevalence continues to increase in low and middle income countries toward epidemic proportions (Roth et al., 2017; Yusuf et al., 2014). Effective tailored strategies for the prevention and treatment depends on thorough understanding of the mechanisms involved in specific populations. The rapid evolution of genomic and personalised precision medicine offers unprecedented opportunities in this regard (Manolio et al., 2009; O’Donnell & Nabel, 2011). These, however, are critically dependent on defining the genetic landscape of different populations, their individuals and the relation to their dynamic phenotype (Lau & Wu, 2018; Leopold & Loscalzo, 2018). In-depth information is lacking in populations which need it most and yet continue to be grossly understudied (Need & Goldstein, 2009; Bustamante, Burchard, & De la Vega, 2011; Owolabi et al., 2014; Popejoy & Fullerton, 2016; Keates, Mocumbi, Ntsekhe, Sliwa, & Stewart, 2017). Notable initiatives and studies have recently begun in Africa and the Middle East and North Africa (MENA) region aiming at data collection and harmonization, paving the way for large scale genomic studies (Scott et al., 2016, Owolabi et al., 2018). To directly address these issues, we are recruiting 1000 Egyptian healthy volunteers (EHVol) from the general population. These individuals are fully phenotyped with respect to cardiovascular health. To date, 724 have been recruited. Here, we describe the protocol of the study and examine background genetic variation in genes previously shown to be involved in inherited cardiac conditions (ICCs), with a special focus on hypertrophic cardiomyopathy (HCM) and dilated cardiomyopathy (DCM). Here we report the data from a pilot cohort of 391 EHVols who were phenotyped and sequenced across a panel of 174 genes involved in ICCs. To our knowledge, this study represents the first of its kind in the region to integrate high coverage sequencing data from a clinically phenotyped cohort.

## METHODS

### Study Protocol and Data collection

This is a population-based study that aims to recruit 1000 Egyptian healthy volunteers (EHVols) from across Egypt. All participants provided informed consent, which was approved by the research ethics committee (20160401MYFAHC_HVOL). Here, we report data from an initial cohort of 391 volunteers recruited from December 2015 to June 2018 via national advertisement (brochures, flyers, public events) and assessed at the Aswan Heart Centre (AHC). A paper-based questionnaire was conducted to gather information regarding demographics, health status, smoking and drinking habits, past medical and surgical history, family history, medication and for the identification of consanguineous marriage (defined as self-reported first-cousin marriage). Individuals were excluded if they met any of the following criteria: <18 years of age, non-Egyptian nationality, pregnancy, presentation with known cardiovascular or collagen vascular disease, communication difficulties or contraindication to cardiac magnetic resonance (CMR) (Figure 1). Data was entered into a Research Electronic Data Capture (REDCap) database; and was stored, along with all appropriate documentation, on an access-controlled server (Harris et al., 2009).

**Figure 1.**
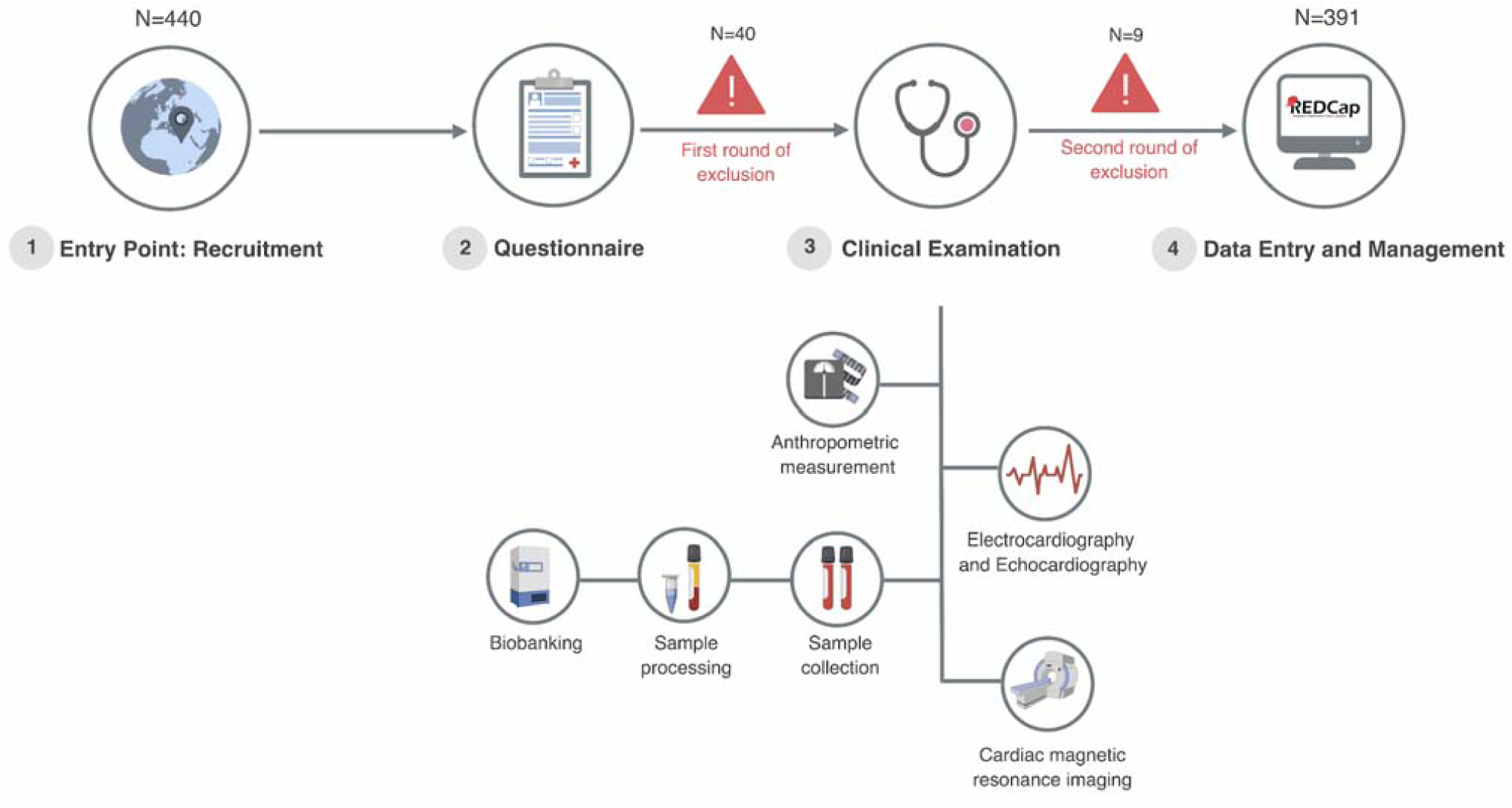
Workflow of EHVol study. Study participants (1) were recruited from the general population via announcements (brochures, flyers, public events); (2) completed a questionnaire on demographic data, family and clinical history; (3) underwent detailed cardiovascular phenotyping and blood sampling. (4) All data were recorded and managed on a local REDCap database. *Rounds of exclusion*: First round of exclusion is based on the basis of demographic and general health questionnaire as described under ‘study protocol and data collection’. A second round of exclusion was based on detailed cardiovascular phenotyping as described under ‘cardiovascular phenotyping’ in the methods section.

### Cardiovascular Phenotyping

All individuals underwent detailed cardiovascular phenotyping including clinical examination, 12-lead electrocardiogram and CMR (Figure1). CMR was performed with a 1.5 T scanner (Siemens Magnetom Aera, Erlangen, Germany) using retrospective ECG triggering to capture the heart during the cardiac cycle. Steady State Free Precession (SSFP) end expiratory breath-hold cine images were acquired in the short axis orientation covering the whole heart. Standard parameters were repetition, time/echo time 3.6/1.8ms; sense factor 2, flip angle, 60°; section thickness, 8 mm; no slice gap, matrix, 160□×□256; field of view, 300 mm; pixel size, 1.6□×□1.6 mm; number of phases 30 and phase percentage 67%. For future comparison with specific disease based sub-cohorts, phase contrast images were acquired at different aortic and levels for flow mapping. At the same acquisition levels 4D flow was performed to assess flow patterns. T1 mapping was performed on base, mid and apical heart levels for fibrosis assessment. 3D Tagging acquisition was done at base, mid and apical levels for strain assessment. Detailed structural and functional analysis on the CMR acquisitions was performed retrospectively using dedicated post processing and in-house software. Following phenotyping, a second round of exclusions, on the basis of specific cardiovascular diagnostic criteria, was applied (Table 1). Isolated apical noncompaction with normal ECG and normal CMR was not excluded.

**Table 1.**
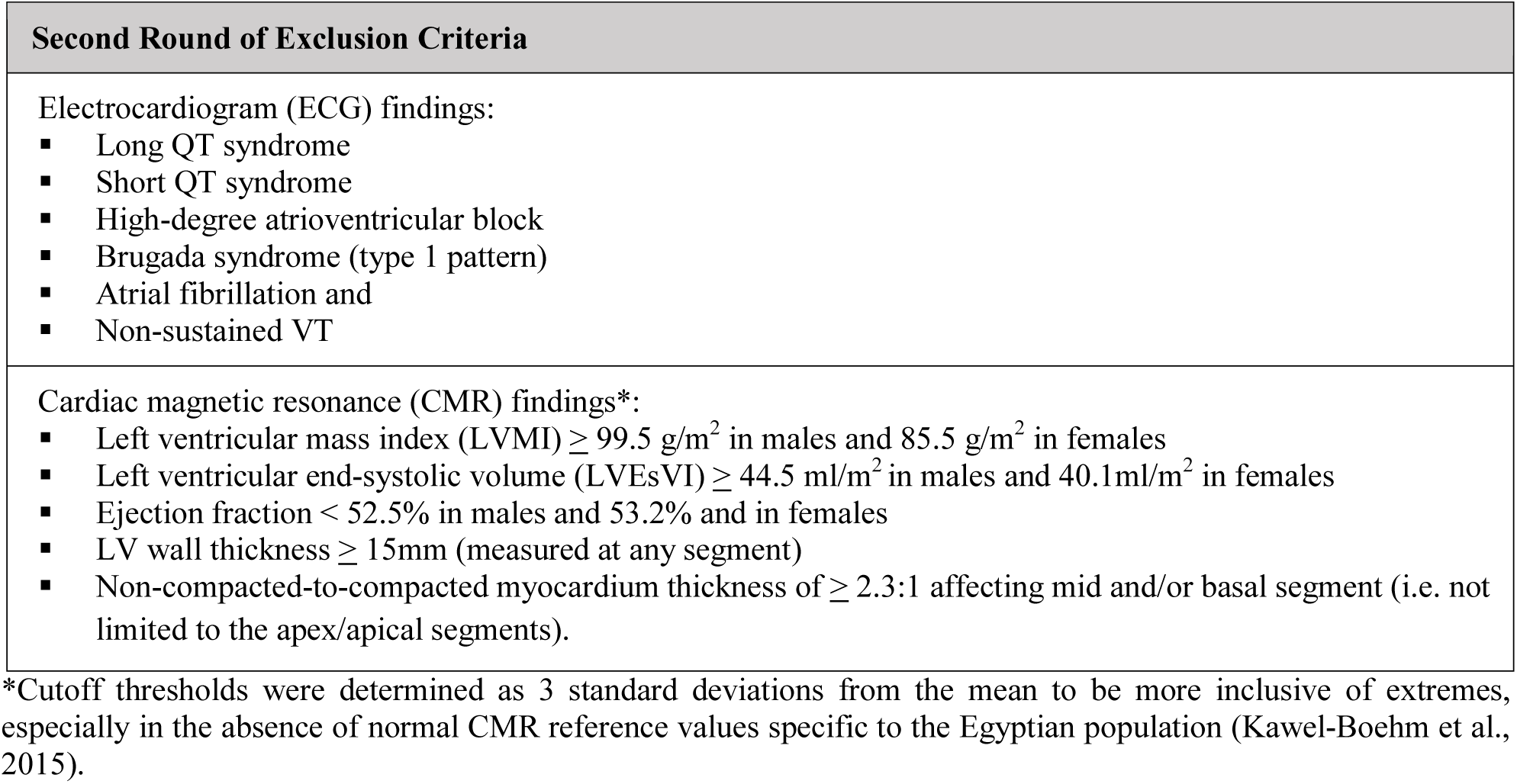
Second Round of Exclusion Criteria.

### Sample Collection and Biobanking

20 ml whole venous blood were withdrawn from each participant for laboratory testing (Hemoglobin A1c (HbA1c) and Troponin I), serum/plasma isolation and DNA extraction. For DNA extraction blood samples were transferred to K3EDTA tubes to avoid clotting. Blood samples were stored at 4oC (max. 5 days) prior to DNA extraction. DNA was extracted using Wizard® Genomic DNA Purification Kit (Promega, Catalog No. A1620) according to manufacturer’s instructions. Concentration of 1μl DNA sample was determined using the NanoDrop 2000 (Thermo Scientific) spectrophotometer. 260/280 and 260/230 nm ratios were used to assess the DNA quality. All samples were stored centrally in the AHC Biobank.

### Next-Generation Sequencing (NGS)

EHVols were sequenced with the Illumina Miseq and Nextseq platforms using the Trusight Cardio Sequencing Kit (Illumina, Catalog No. FC-141-1010 (MiSeq) and FC-141-1011 (NextSeq) comprising 174 genes with reported roles in ICCs (Pua et al., 2016). Sequencing was performed following the manufacturer’s protocol. The concentration and quality of the DNA libraries were evaluated using Qubit (Invitrogen) and TapeStation 4200 (Agilent Technologies)

### Bioinformatics Pipeline

Raw data was subject to quality control using FastQC v0.10.1 (Simon Andrews, 2010) and low-quality reads were trimmed via prinseq-lite v0.20.4 (Schmieder & Edwards, 2011). Trimmed reads were then mapped to hg19 (Kuhn et al., 2009) using the Burrows-Wheeler Aligner (BWA) v0.7.10-r789 (Li & Durbin, 2009). After alignment, removal of duplicate reads was performed using picard v1.117 (“Picard Tools,” n.d.). GATK v3.2-2-gec30cee (McKenna et al., 2010) was then used for re-alignment of insertions or deletions (indels) as well as base quality score recalibration. Variant calling was performed using GATK’s HaplotypeCaller. Joint genotyping is performed with GATK v4.0.8.1 GenotypeGVCFs and hard filters were applied based on GATK best practises workflow for germline short variant discovery (McKenna et al., 2010). Only variants that are marked as “PASS”,with a quality of depth score (QD) ≥ 4 and an allelic balance (AB) ≥ 0.20 were retained. We attained high coverage of the target region (>99%) at ≥ 20× read depth (Supplementary Figure S1). The Ensembl Variant Effect Predictor (VEP) (McLaren et al., 2016) was used for variant annotation.

### Data Analysis

#### Analysis of background genetic variation in selected CM genes

To examine the background genetic variation among genes involved in inherited CMs; specifically DCM and HCM, we selected genes that were recently validated (Supplementary Table S1). The following DCM genes were analysed: *BAG3, DSP, LMNA, MYH7, RBM20, SCN5A, TCAP, TNNC1, TNNT2, TPM1, TTN* and *VCL (*Walsh et al., 2017). Genes with definitive evidence of HCM association, such as *MYBPC3, MYH7, TNNT2, TNNI3, TPM1, ACTC1, MYL2, MYL3* and *PLN* were studied (Ingles et al., 2019). In addition, syndromic genes definitively associated with isolated left ventricular hypertrophy (LVH), such as *CACNA1C, DES, FHL1, GLA, LAMP2, PRKAG2, PTPN11, RAF1* and *TTR* were analysed (Ingles et al., 2019).

#### Comparison of genetic variation between EHVol and gnomAD controls

The Genome Aggregation Database (gnomAD) is the largest population data set to date. It comprises genetic data from 125,748 and 15,708 unrelated individuals sequenced by whole exome sequencing (WES) and whole genome sequencing (WGS), respectively (Karczewski et al., 2019). We downloaded WES data from the gnomAD database (https://gnomad.broadinstitute.org; version v2.1.1; Karczewski et al., 2019). Only high quality (‘PASS’) variants were included in the analysis. Genetic variation in CM genes was compared between the EHVol and gnomAD controls. Characterized variant types included loss-of-function (LoF) (i.e. frameshift, splice acceptor, splice donor, nonsense), missense (i.e. missense, inframe deletion, inframe insertion), synonymous and other (3’ and 5’ UTRs, intronic, splice region…etc).

#### Comparison of rare variation between EHVol and Caucasian HVOL (CHVol) controls

A cohort of 1,028 Caucasian healthy volunteers (CHVols) also sequenced using the Trusight Cardio Sequencing Kit for ICC, was analysed (Schafer et al., 2017; Pua et al., 2016). The CHVols were recruited prospectively via advertisement for the UK Digital Heart Project at Imperial College London. All volunteers underwent CMR to confirm the absence of cardiac disease. The frequency of rare variation in the selected CM genes was compared between the EHVol and CHVol cohorts. The threshold maximum credible population allele frequencies (AF) were defined as <=8.4×10-5 and <=4.0×10-5 for DCM and HCM, respectively. Variants were defined as rare if the filtering allele frequency (FAF) was less than these thresholds across all gnomAD populations (popmax FAF) (Whiffin et al. 2017). The frequency of rare variants per gene in the EHVol/CHVol cohorts was calculated by counting the number of rare variants per gene and dividing this by the cohort size (EHVols: n=391, CHVols: n=1,028).

## RESULTS

### Characteristics of the EHVol Study Population

Of the 440 recruited individuals, 40 met the first round of exclusion criteria and were therefore excluded. The remaining individuals underwent CMR (n=400) and ECG (n=349) screening (table 2). Based on the second round of exclusion described above, nine individuals were excluded from the cohort (Supplementary Table S2). A final cohort of 391 EHVols were sequenced. The baseline characteristics of the study population are summarized in table 3. The study population comprised of 166 females (42.5%) and 225 males (57.5%). The mean age (years) was 33.2 (SD 9.5).

**Table 2:** General cardiac characteristics of the phenotyped individuals (n=400.)

| Cardiac characteristics | Mean ± Standard deviation |
| --- | --- |
| **ECG variables** |  |
| PR interval (ms) | 147 ± 22 |
| QRS duration (ms) | 91 ± 47 |
| QTc interval (ms) | 400 ± 45 |
| Abnormal ECG*, N( %) | 19 (4.8) |
| **CMR Variables** |  |
| LVEF (%) | 63 ± 5.7 |
| LVMI (g/m<sup>2</sup>) | 47 ± 14 |
| LVESVI (ml/m<sup>2</sup>) | 28 ± 7.8 |
| LVEDVI (ml/m<sup>2</sup>) | 75 ± 12.2 |
| Abnormal CMR*, N (%) | 11 (2.8) |

\*Abnormality is defined according to the American Board of Internal Medicine (ABIM) coding criteria

**Table 3:**
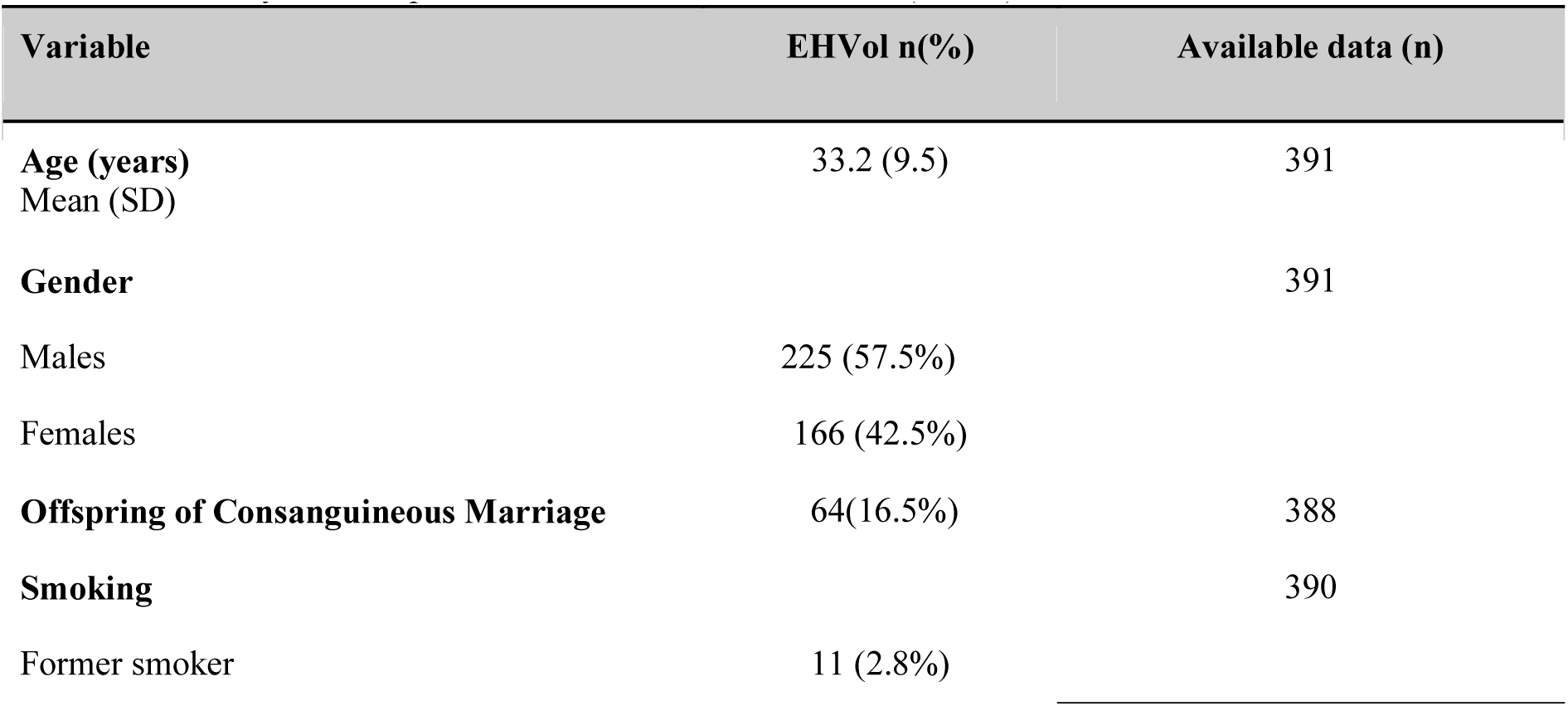
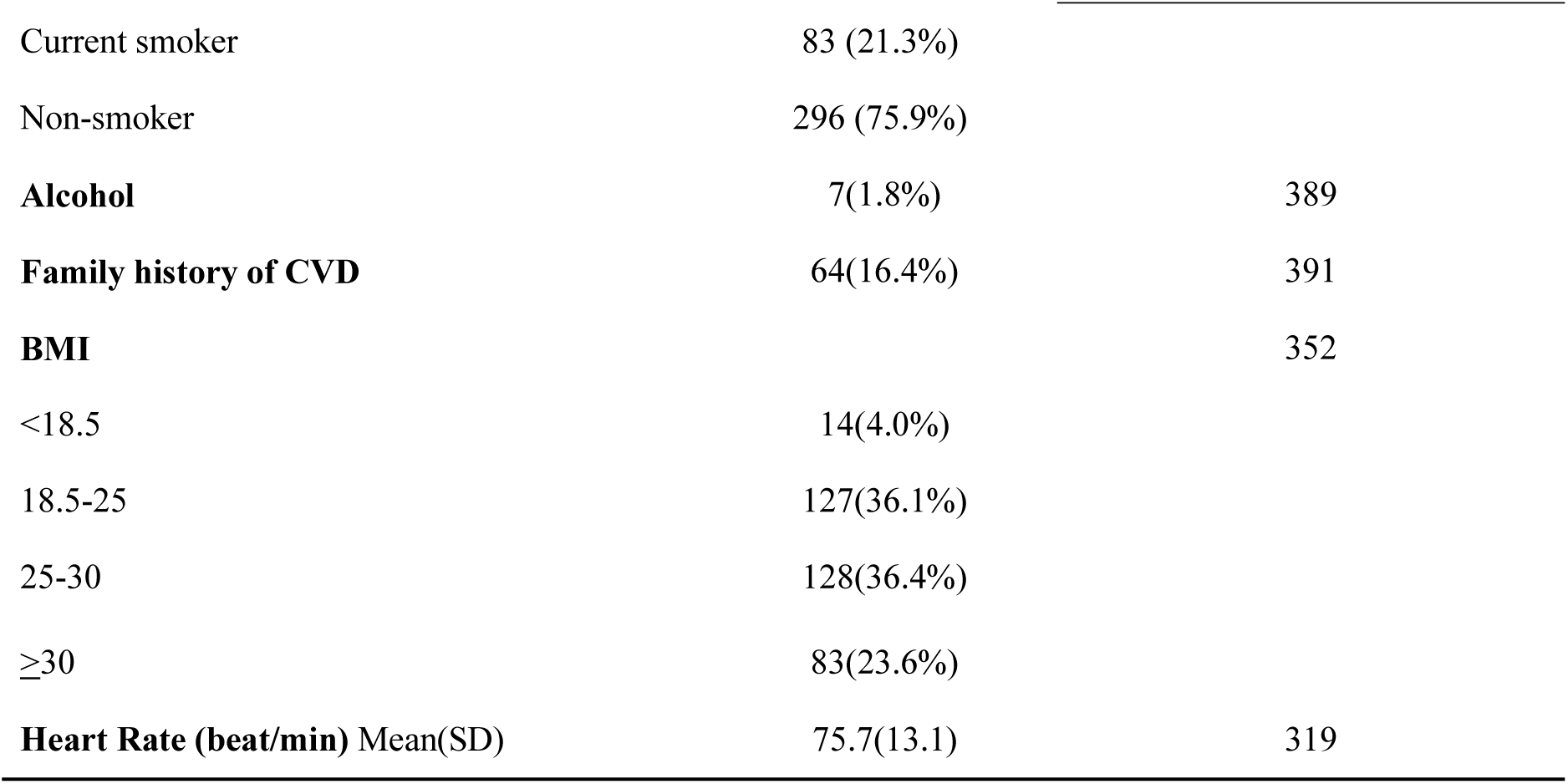
Summary of descriptive characteristics of the EHVols (n=391).

| Variable | EHVol n(%) | Available data (n) |
| --- | --- | --- |
| **Age (years)**<br>Mean (SD) | 33.2 (9.5) | 391 |
| **Gender** |  | 391 |
| Males | 225 (57.5%) |  |
| Females | 166 (42.5%) |  |
| **Offspring of Consanguineous Marriage** | 64(16.5%) | 388 |
| **Smoking** |  | 390 |
| Former smoker | 11 (2.8%) |  |
| Current smoker | 83 (21.3%) |  |
| Non-smoker | 296 (75.9%) |  |
| **Alcohol** | 7(1.8%) | 389 |
| **Family history of CVD** | 64(16.4%) | 391 |
| **BMI** |  | 352 |
| <18.5 | 14(4.0%) |  |
| 18.5-25 | 127(36.1%) |  |
| 25-30 | 128(36.4%) |  |
| ≥30 | 83(23.6%) |  |
| **Heart Rate (beat/min) Mean(SD)** | 75.7(13.1) | 319 |

### Representation of EHVol genetic variation in gnomAD

2,040 CM variants were identified in the EHVol cohort (Supplementary Table S3). In order to assess the representation of these variants in the gnomAD dataset, we plotted the observed allele counts (ACs) in gnomAD against ACs in EHVol for all variants identified in our cohort. These EHVol variants were also binned by AC to report the proportion that was captured in gnomAD (Figure 2A). Of the 2,040 EHVol CM variants, 1544 (75.7%) were captured in gnomAD, whereas 496 (24.3%) were absent from gnomAD (Table 4). The majority of non-gnomAD variants (n=335) were captured in AC bin 1. The remaining non-gnomAD variants constituted <10% of each AC bin (Figure 2A). Non-gnomAD variants were predominantly missense (29.8%) and “other” (56.5%) (Figure 2B). Out of the 496 non-gnomAD variants identified in the EHVol cohort, 11.3% were present in the Great Middle Eastern Variome (GME) (Scott et al., 2016). The proportion of EHVol non-gnomAD variants was significantly (Fisher’s exact test p<1.183e-07) higher than that of the CHVol cohort (24.3% vs 18.3% respectively) (Table 4).

**Table 4:** Proportion of non-gnomAD variants in EHVol and CHVol cohorts.

| Cohort | Cohort Size | Total no. of variants in CM & syndromic genes | Total no. of non-gnomAD variants<sup>†</sup> | Prop. of non-gnomAD variants |
| --- | --- | --- | --- | --- |
| EHVol | 391 | 2,040 | 496 | 24.3%* |
| CHVol | 1,028 | 3,592 | 658 | 18.3%* |

<sup>†</sup> Only 'PASS' variants in gnomAD were included in the analysis
\*=Fisher's exact test p-value < 1.183e-07

**Figure 2.**
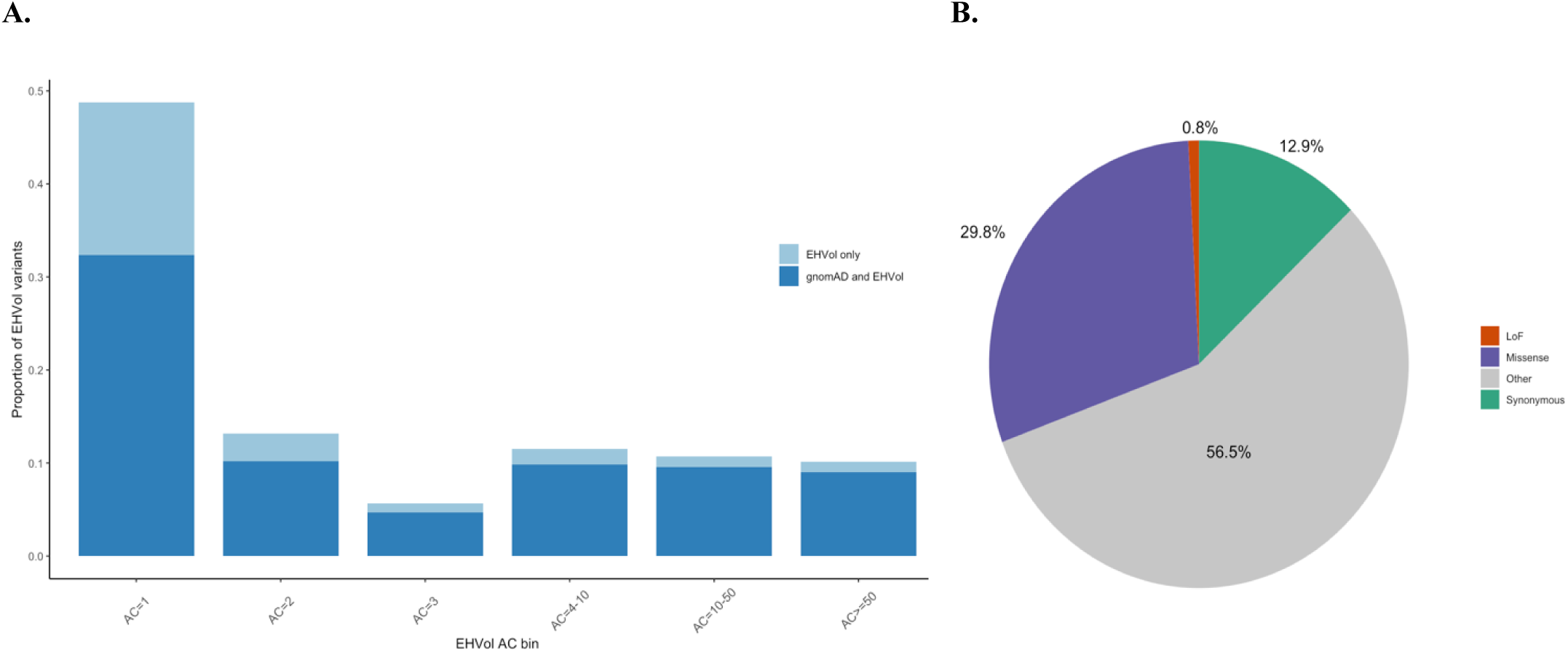
(A) Bar graph of the observed ACs (binned) in all CM variants identified in the EHVol cohort. The proportion of variants that were captured in gnomAD (all populations) is shown in dark blue and the proportion of variants that were only captured in our cohort is represented in light blue. (B) Pie chart showing the distribution of non-gnomAD variants by variant type.

### High frequency of rare variation in the EHVol cohort

The proportion of EHVol and CHVol controls with rare variants in DCM (popmax FAF <=8.4×10-5) and HCM (popmax FAF <=4.0×10-5) genes was calculated (Figure 3). In both cohorts, *TTN, DSP, RBM20, MYH7* and *SCN5A* accounted for the majority of rare variation in DCM genes (Figure 3A). *MYBPC3, MYH7* and *CACNA1C* accounted for the majority of rare variation in HCM genes in both cohorts (Figure 3B). Overall, the frequency of rare variants was higher among EHVols compared to CHVols. The proportion of controls with LoF variants in CM genes was almost the same in the EHVol (2.3%) and CHVol (2.33%) cohorts.

**Figure 3.**
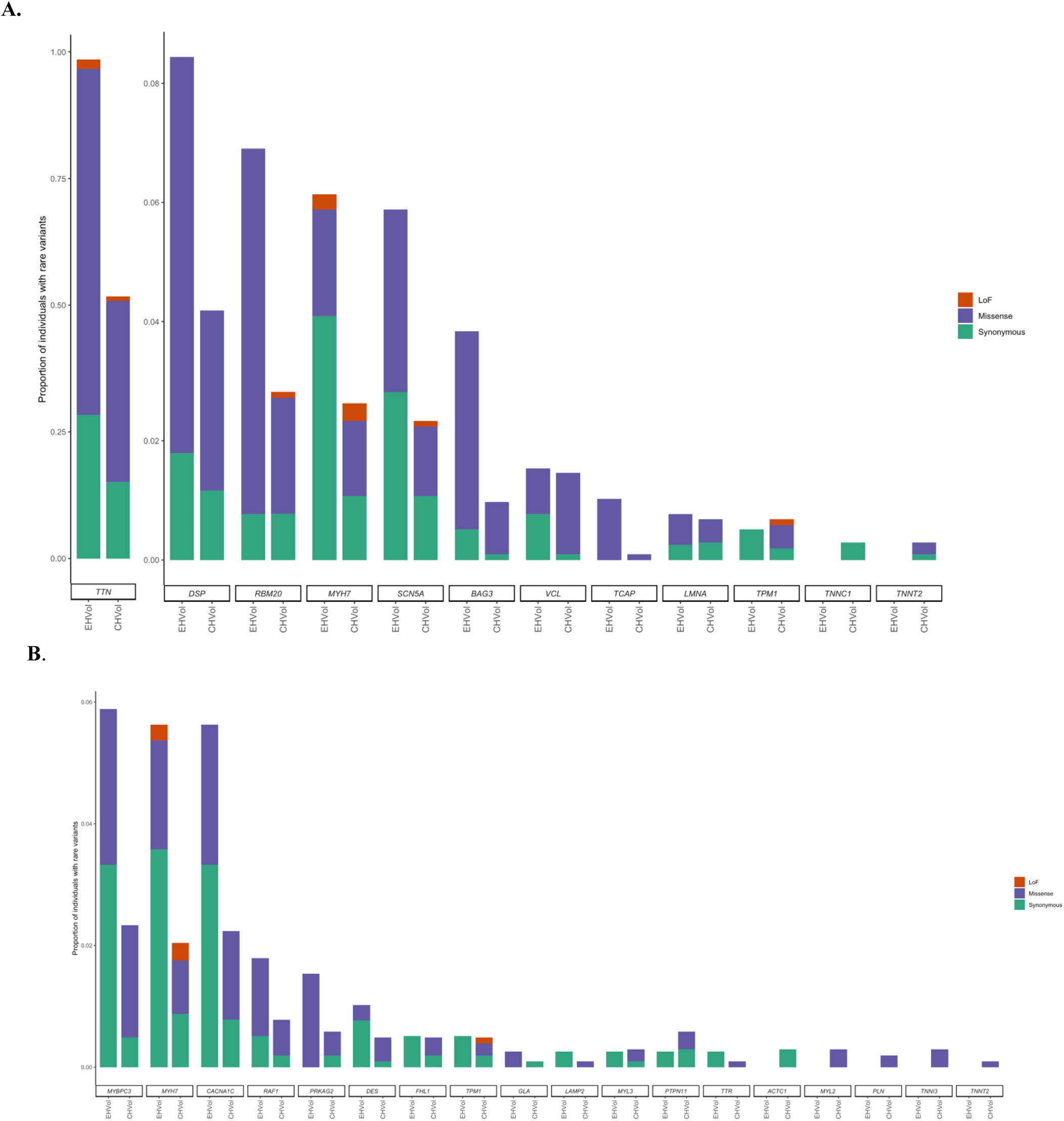
Rare variation in CM genes in EHVol and CHVol controls. Bar charts represent the proportion of individuals with rare variants in (A) DCM and (B) HCM and LVH syndromic genes. The first and second bars represent the CHVol and EHVol cohorts, respectively. Variants are collated by variant type (LoF, Missense and Synonymous).

### Comparison of the distribution of rare *TTN* and *MYH7* variants between the different control cohorts

We investigated the distribution of variants in *TTN* and *MYH7*, as they accounted for relatively high frequencies of rare variation among DCM and HCM genes, respectively. We restricted the analysis to rare LoF variants in *TTN* and missense and LoF variants in *MYH7* (Figure 4), as they may be putative pathogenic. *TTN* consists of four regions, Z-disk, I-, A- and M-bands, each of which has a distinct function (Roberts et al., 2015). Of the four *TTN* regions, the I-band has the lowest expression level in the myocardium (i.e. percentage spliced in (PSI) <0.9), as I-band exons are variably spliced in different isoforms (Roberts et al., 2015). Across all cohorts, the majority of variants were located in exons that are not constitutively expressed in the myocardium (Figure 4A). One LoF variant (p.Ser34842ProfsTer9) and 6 variants (p.Glu35478Ter, p.Gln16235Ter, p.Gln15575Ter, c.44015-1G>T, p.Pro4353GlnfsTer14 and p.Gln3243Ter) identified in the EHVol and CHVol cohorts, respectively, resided in cardiac constitutive exons, which would have been interpreted as “likely pathogenic” (LP) were they identified in individuals presenting with DCM (Roberts et al., 2015). The LoF variant p.Ser34842ProfsTer9 identified in the EHVol cohort, is absent among gnomAD as well as CHVol controls. The EHVol carrying the LoF variant did not show a clinical DCM phenotype but ECG analysis revealed sinus tachycardia.

**Figure 4:**
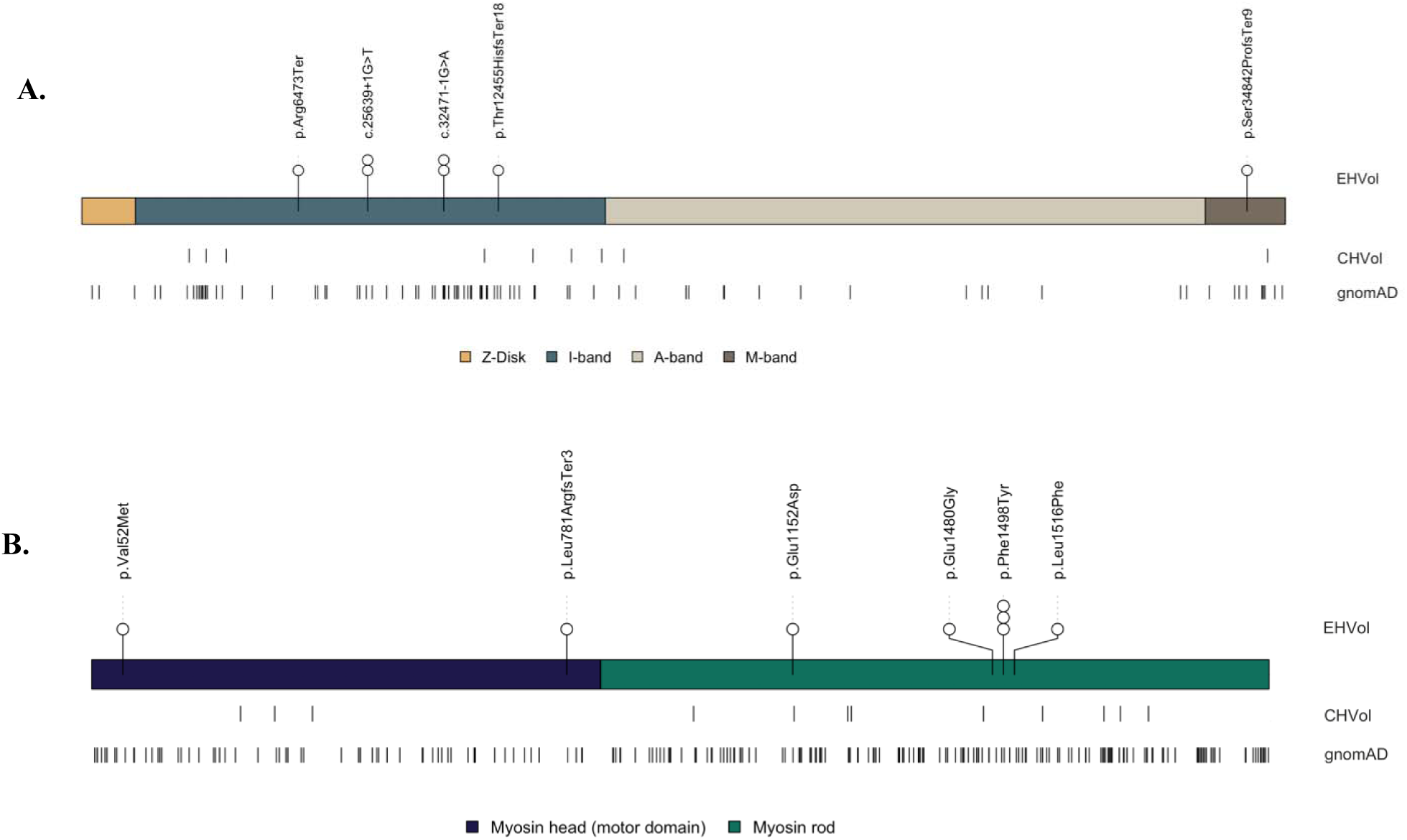
Distribution of rare (A) *TTN* LoF and (B) *MYH7* missense and LoF variants in the EHVol, CHVol and gnomAD cohorts. Variant distribution is shown relative to a schematic representation of the *TTN* protein, with sarcomere regions delimited and the *MYH7* protein with myosin domains delimited. The number of circles represent the number of individuals carrying the rare variant.

Across all populations, the majority of rare variants in *MYH7* (popmax FAF <=4.0×10-5) were located in the rod domain (EHVol 4 (66.7%); CHVol 9 (75%); gnomAD 163 (71.5%)) (Figure 4B).

## DISCUSSION

The availability of large-scale population genetic databases, such as gnomAD, has opened up new possibilities in identifying putative disease-causing variants. Current data sets, however, do not adequately represent genetic variation in the MENA region, which hinders accurate variant interpretation in the respective populations. Several recent studies address this shortcoming. Scott et al., for instance, studied rare genetic variation of the Great Middle Eastern region, including North East- and North West Africa (Scott et al., 2016). However, individuals recruited in this study were from the general population (self-reported healthy volunteers) and thus may not necessarily be free of disease. To our knowledge, our study represents the first of its kind in the region to include high coverage sequencing data from a clinically phenotyped healthy cohort. We evaluated 440 self-declared healthy individuals of whom 49 (11%) were excluded for not meeting the first round inclusion criteria (9%) or showed cardiac abnormalities (2%). The remaining healthy individuals, the EHVols (n=391), were genetically characterised using the ICC gene panel. Our analyses revealed that 24.3% of EHVol variants were not captured in gnomAD, the most widely used reference dataset at present. Of these non-gnomAD variants 11.3% were captured in the GME database, confirming the current underrepresentation of genetic variation in current large-scale datasets.

In addition, we sought to compare genetic variation between our EHVols and an ethnically-distinct population that is generally represented in gnomAD, the CHVol cohort. Our analysis showed that the proportion of non-gnomad variants identified in the EHVol cohort was significantly higher compared to the CHVol cohort. Furthermore, we examined the frequency of putative disease-causing variants in the EHVol and CHVol cohorts across the CM genes. The analysis showed that the proportion of controls with rare variation in the majority of DCM genes was higher in the EHVol cohort compared to CHVols. Rare variation in *MYH7* and *MYBPC3*, the key genetic contributors to HCM, was also higher in the EHVol cohort.

These findings highlight the importance of studying the prevalence of putative disease-causing variants in a large-scale ethnic-specific cohorts in order to confirm their pathogenicity in their respective populations. For example, a novel, non-gnomAD LoF *TTN* variant (p.Ser34842ProfsTer9) was identified in a 33-year old EHVol. This variant lies in the M-band, a constitutively expressed exon with a PSI score of >0.9, which affects cardiac remodeling in DCM (Roberts et al., 2015; LeWinter & Granzier, 2013). Had this LoF variant been identified in a well-defined DCM cohort, it would have been interpreted as a “likely pathogenic” (LP). Thus, it would be valuable to follow-up on EHVols carrying putative disease-causing variants to confirm their corresponding phenotypic manifestations. Integrating information from our EHVol cohort into genomic datasets and variant classification support tools, such as CardioClassifier, may affect the classification of detected variants.

Our analysis has provided preliminary insights into genetic variation of the underrepresented Egyptian population. This study constitutes an initial milestone towards developing a large-scale dataset comprising healthy EHVols from across the country. The development of an ancestry-specific genetic and phenotypic database will circumvent current issues with variant interpretation, deepen our understanding of African diversity and provide an opportunity for variant discovery. The expanded dataset will also directly aid in distinguishing between incidental and medically actionable variants and thus enhance diagnostic and treatment strategies. Beyond the ethnicity-specific aspects, this study provides valuable molecular and phenotypic data and a regional biobank.

A limitation of this study is that allele frequency thresholds used in our analysis may not accurately define rare variants in CM genes in our population. That is because these thresholds were pre-computed by Whiffin et al., 2017 based on genetic findings from a Caucasian CM cohort. Also, future cohort studies might show that there are pathogenic variants above this threshold e.g. due to a founder. However, using different frequency thresholds in different populations would introduce an artifactual difference in rare variant burdens, as it would vary each population’s definition of rare. A suitable future approach is to adjust filtering allele frequencies (FAF) whenever data from new populations are integrated into a large data set.

## ACKNOWLEDGMENTS

We thank the Egyptian volunteers for their generous contribution to the study. We also thank Ms. Omayma Alm El Din for her dedication and diligence. We would like to thank the Genome Aggregation Database; and all the contributing groups that provided exome variant data for comparison (full list can be found at: http://gnomad.broadinstitute.org/about).

This study was supported by the Science and Technology Development Fund (STDF) government grant.

