## Supplemental Figure for "Genomics of Egyptian Healthy Volunteers: The EHVol Study"

**B**

**A**

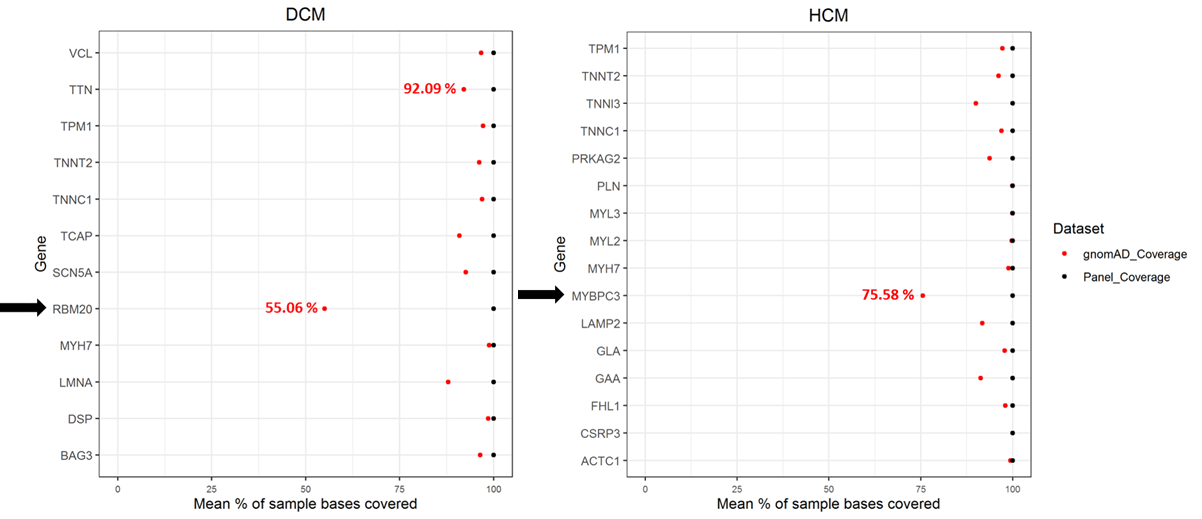

**Figure S1: Comparison of sequencing coverage in DCM/HCM genes between EHVol (ICC Panel) and gnomAD version 2.1 data (WES).**

*Figure S1: Sequencing coverage differences between EHVol and gnomAD for: (a) the 12 selected DCM genes and (b) the 16 selected HCM genes. The x-axis represents the percentage of bases covered for each gene at 20X depth of coverage. Mean percentage of sample bases covered for RBM20, TTN and MYBPC3 are indicated in red.*
